# Perceived Risk and Barriers to Open and Responsible Research Across Fifteen UK Universities

**DOI:** 10.64898/2026.09.04.746981

**Authors:** Noémie Aubert Bonn, Andrew J. Stewart, Lukas Hughes-Noehrer

## Abstract

Open research practices are increasingly promoted to improve research transparency, reproducibility, accessibility, and societal impact. Despite growing support from research funders, institutions, and policy initiatives, adoption remains uneven across disciplines and research communities. This study examined perceived risks and barriers associated with 14 FORRT Guideline areas using qualitative responses from the UK Reproducibility Network Open and Transparent Research Practices Survey (N = 2,567), conducted across 15 UK higher education institutions. Free-text responses describing risks and barriers were analysed using inductive thematic analysis.

A total of 3,951 relevant comments generated 3,433 coded references to barriers and risks. Three interconnected clusters emerged. Individual concerns included lack of motivation, fear of losing intellectual credit, and concerns about exposing mistakes and criticism. Systemic and institutional barriers included lack of time and resources, inadequate infrastructure, insufficient training and support, lack of incentives and recognition, unclear guidance, disciplinary and methodological challenges, and tensions between open practices and intellectual property requirements. Ethical and quality-related concerns included risks to participant confidentiality and privacy, challenges associated with sensitive data, concerns about inappropriate application of open research practices across different research traditions, perceived impacts on research quality and innovation, and potential effects on public trust in research. Lack of time and resources was the most frequently reported barrier. Across responses, barriers were commonly described as interdependent, with shortcomings in funding, infrastructure, institutional support, incentives, and training reinforcing one another.

Respondents largely supported the principles underpinning open research but highlighted substantial practical, professional, and ethical challenges to implementation. These findings suggest that increasing open research adoption requires more than policy mandates or awareness-raising activities. Sustainable uptake will depend on aligned incentives, adequate infrastructure and support, recognition of methodological diversity, and approaches that enable openness to be implemented responsibly across different research contexts.

## 1 Introduction

Open research practices are increasingly recognised as a foundation for transparent, accessible, and rigorous research that maximises its societal value and impact. Across disciplines, concerns regarding research quality, reproducibility, selective reporting, publication bias, and limited access to research outputs have prompted calls for greater openness throughout the research lifecycle [1, 2].

Growing evidence suggests that open research practices can improve transparency, facilitate collaboration, enhance reproducibility, accelerate knowledge dissemination, and increase opportunities for innovation and societal impact [3, 4]. Funders such as UK Research and Innovation (UKRI), Wellcome Trust, the National Institutes of Health (NIH), and the European Commission have embedded open research expectations within funding policies, whilst universities are increasingly developing institutional strategies to support open and reproducible research. At the same time, initiatives such as the UK Reproducibility Network (UKRN) have sought to build infrastructure, training, communities of practice, and institutional capacity to facilitate cultural change across the UK research landscape [5].

Despite widespread endorsement of open research principles, evidence consistently demonstrates that implementation remains uneven across disciplines, institutions, and career stages. Researchers frequently report barriers associated with limited time, insufficient resources, inadequate training, lack of incentives, uncertainty regarding ethical and legal requirements, and concerns about potential adverse impacts on career progression [6–8]. Additional concerns have been raised regarding the applicability of particular open research practices to qualitative, exploratory, creative, or sensitive research contexts, suggesting adoption challenges may differ substantially between research communities [9, 10].

The successful implementation of open and responsible research therefore depends not only on policy mandates, but also on understanding how the research community perceives the risks and barriers associated with these practices. Although previous studies have explored attitudes towards individual elements of open research, such as data sharing, pre-registration, or open access publishing, fewer investigations have examined barriers systematically across a broad range of open research practices and institutional contexts. Such evidence is essential for developing targeted interventions, training programmes, infrastructure, and policy approaches that address the practical realities faced by research communities and to support the effective and sustainable adoption of open and responsible research practices, consistent with international recommendations for capacity building and cultural change in research environments [11].

This study draws on data from the UKRN Open and Transparent Research Practices (OTRP) Survey [12], conducted across fifteen UK Higher Education Institutions (HEIs). Using qualitative thematic analysis of researchers’ free-text responses, we examine perceived risks and barriers associated with fourteen open research practice areas. Our aim is not only to identify the specific challenges encountered, but also to understand how these concerns cluster across individual, institutional, and ethical dimensions. By providing a comprehensive overview of researchers’ perspectives across a diverse UK research landscape, this study seeks to inform future efforts to support the effective, equitable, and sustainable adoption of open and responsible research practices.

## 2 Methods

We used openly available data from the UKRN OTRP. Details on the methodology used for the survey is available in the dedicated data descriptor [12].

The survey was built to capture ongoing awareness and involvement in Open Research practices in UK HEIs, as well as to obtain an overview of the support and training needs that could help improve uptake of Open Research practices. The survey was shared with research-active staff in 15 HEIs in the UK between December 2022 and April 2023. Participants were asked to respond to every question in the survey. In total, 2,567 participants took part in the survey. Of these, 1,221 (48%) answered at least half of the survey questions, including 1,081 (42%) who completed the survey.

The survey was divided into 14 topics covering a range of open research practices. The 14 practice areas were selected based on consensus obtained within a committee of UKRN Institutional Leads (ILs) recruited from institutions partaking in the UKRN Open Research Programme [12]. Selected themes were defined according to the terminology used in the Framework for Open and Reproducible Research Training (FORRT) community-sourced glossary [13]. The topics included in the survey are (i) research co-production; (ii) conduct of open research consistent with relevant legal, ethical and regulatory constraints; (iii) transparent qualitative data; (iv) data management; (v) pre-registration of research protocols; (vi) use of open source software; (vii) creation of open source software; (viii) version control of research products; (ix) computational reproducibility of data analysis; (x) sharing of data, code, or other evidence according to the FAIR principles; guidelines for recognising the specific substantive contribution of everyone involved in research projects; declaration of interests; (xiii) publication of preprints; and (xiv) open access publishing. For each topic, the survey assessed respondents’ own practices; perceived importance of the topic; available support, help, and training; and potential risks and barriers to undertaking the practices in their institution. This paper relates to the latter element, analysing qualitative answers to the open-box question *“If this topic involves risks or barriers to you or to your field of study, please briefly explain here”* which was asked for all fourteen topics (question numbers 1.12, 2.12, 3.12, 4.12, 5.12, 6.12, 7.12, 8.12, 9.12, 10.12, 11.12, 12.12, 13.12, and 14.12). Alongside these practice-specific results, we also looked at the final question from the survey which was a multiple choice question asking respondents *“Having considered barriers above, which do you perceive to be as the greatest barrier to the uptake of open research practices in your field?”* Respondents could select a response among 10 different choices, or they could indicate ‘Other’ and specify their responses.

A response to these questions was mandatory in the survey, but respondents could write ‘NA’, ‘No answer’, or other answers which indicated that they had nothing to add on the matter. Only answers which did not imply a non-response were coded. We analysed responses through inductive thematic coding [14] using NVivo [15] to organise answers. The analysis was conducted using openly available data from 24 July 2024. The survey was anonymous and as such, the authors did not have access to identifiable information, despite their role in creating the initial dataset.

Themes were created inductively based on the answers provided, and reorganised at several points during the coding to build a coherent overview of the concerns raised. Individual answers could be coded in single or in multiple themes to provide a complete overview of the themes perceived as risks and barriers for each answer. For example, the answer “It’s mainly to do with support and training, and having wider understanding of what the risks are at the beginning of the research and not at the end which then prevents you from sharing knowledge and mobilising change” would be coded in the themes of ‘Lack of support’, ‘Lack of training’, and ‘Time and resources’ (through the sub-code ‘Requires early thinking’). When comments only contained positive comments addressing the importance of the open research practice, they were classified in a general counter-node to highlight that the comment reflected on the importance, existing support, or already routine open research practice, rather than barriers or risks. At different points in the coding process, the researcher responsible for the coding (NAB) met with co-authors (LH-N, AJS) to discuss the emerging themes, to merge, refine, and group relevant themes, and to identify emerging clusters.

Coded themes and clusters were later arranged in a thematic matrix highlighting the number of instances a specific theme was mentioned as a risk for any given Open Research Practice. Given the qualitative nature of the analysis and the unequal number of responses for each research practice, these numbers should only be used to identify and illustrate recurrent themes, but they should not be used to compare between open research practices.

## 3 Results

### 3.1 Responses analysed

Over the 14 questions posed in the survey, 3951 comments were coded as containing information (i.e., going beyond ‘N/A’, ‘no comments’, or other non-answers) into 4,492 instances of code. From these, 3,433 codes were placed into risks and barriers. In addition, 752 codes were listed as counter nodes (i.e., mentioning how important or how well achieved an open research practice is rather than highlighting a risk or a barrier) and 307 codes were marked as unclear or irrelevant (e.g., comments on the survey or answer that was unclear of irrelevant such as *“I did this so long ago I cannot say”*) and were not included into further analyses. Three clusters of barriers and risks emerged from the diverse themes coded, namely (A) individual concerns, (B) systemic and institutional constraints, and (C) ethical and quality concerns. In the next sections, we will look at each of these clusters and describe the themes they contain, highlighting examples of concerns raised by survey respondents. An overview of the number of comments coded for each theme in each open research practice is available in Table 1, and an overview of the overall distribution of themes is presented in Figure 1. A few of the quotes included are minimally edited to correct for typos or grammatical error when the original respondent word choice was obvious (e.g., “authroship” was changed to ‘authorship’, ‘barrriers’ was changed to ‘barriers’, etc.), but the majority of quotes are reported without any changes. A dataset containing all responses analysed and the themes and clusters they are associated with is available on Figshare [16].

**Table 1.**
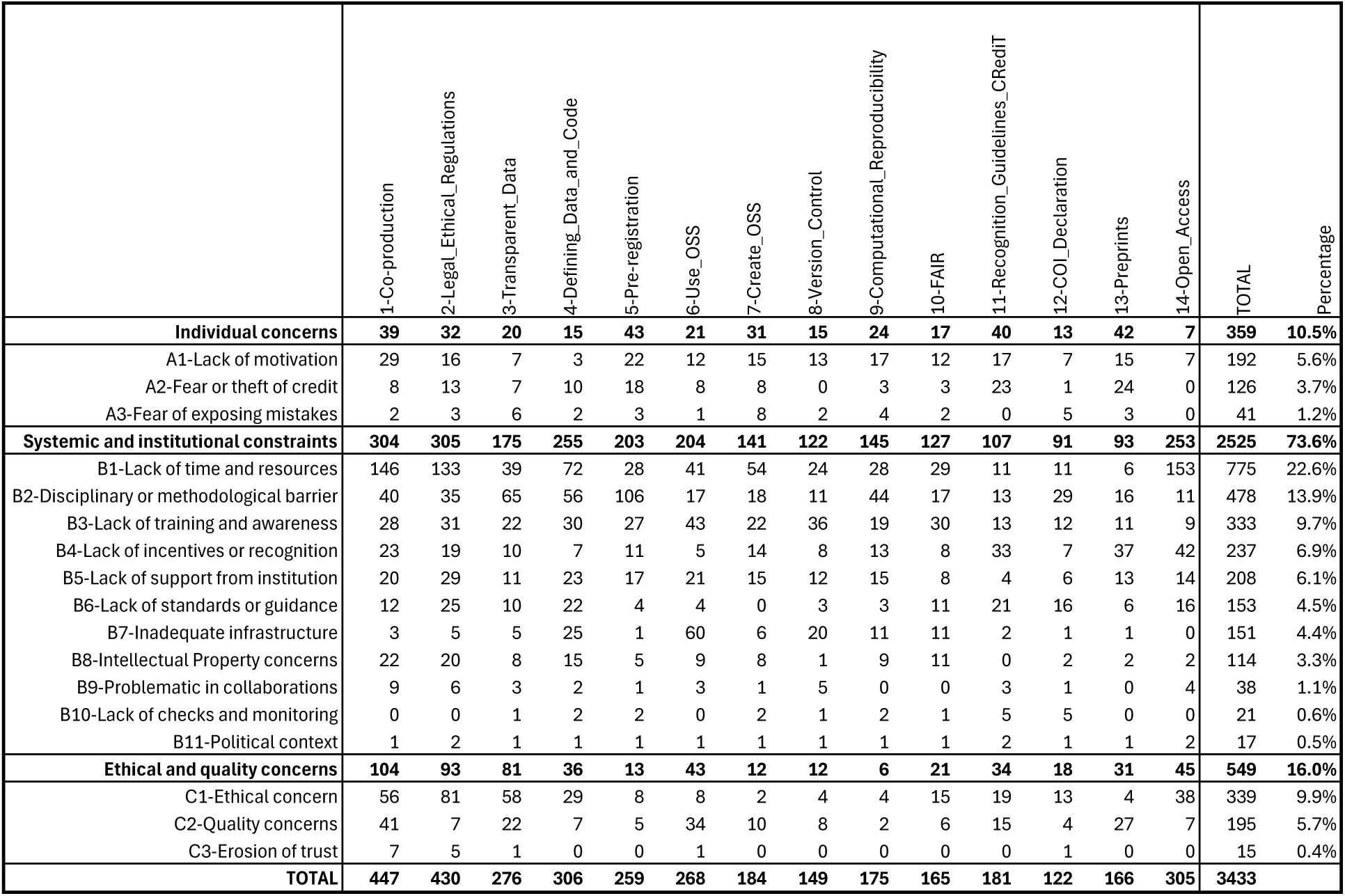
Number of comments coded for each theme in each open and responsible practice.

### 3.2 Cluster A - Individual concerns

A few of the themes identified through the responses were clustered as individual concerns, namely issues related to individual factors, motivation, and personal concerns towards open research practices. Among these, respondents expressed lacking motivation to undertake open research, worrying that open research practices could lead to others stealing or profiting from their work or that they could make them vulnerable by exposing errors and issues with their work.

**Fig 1.**
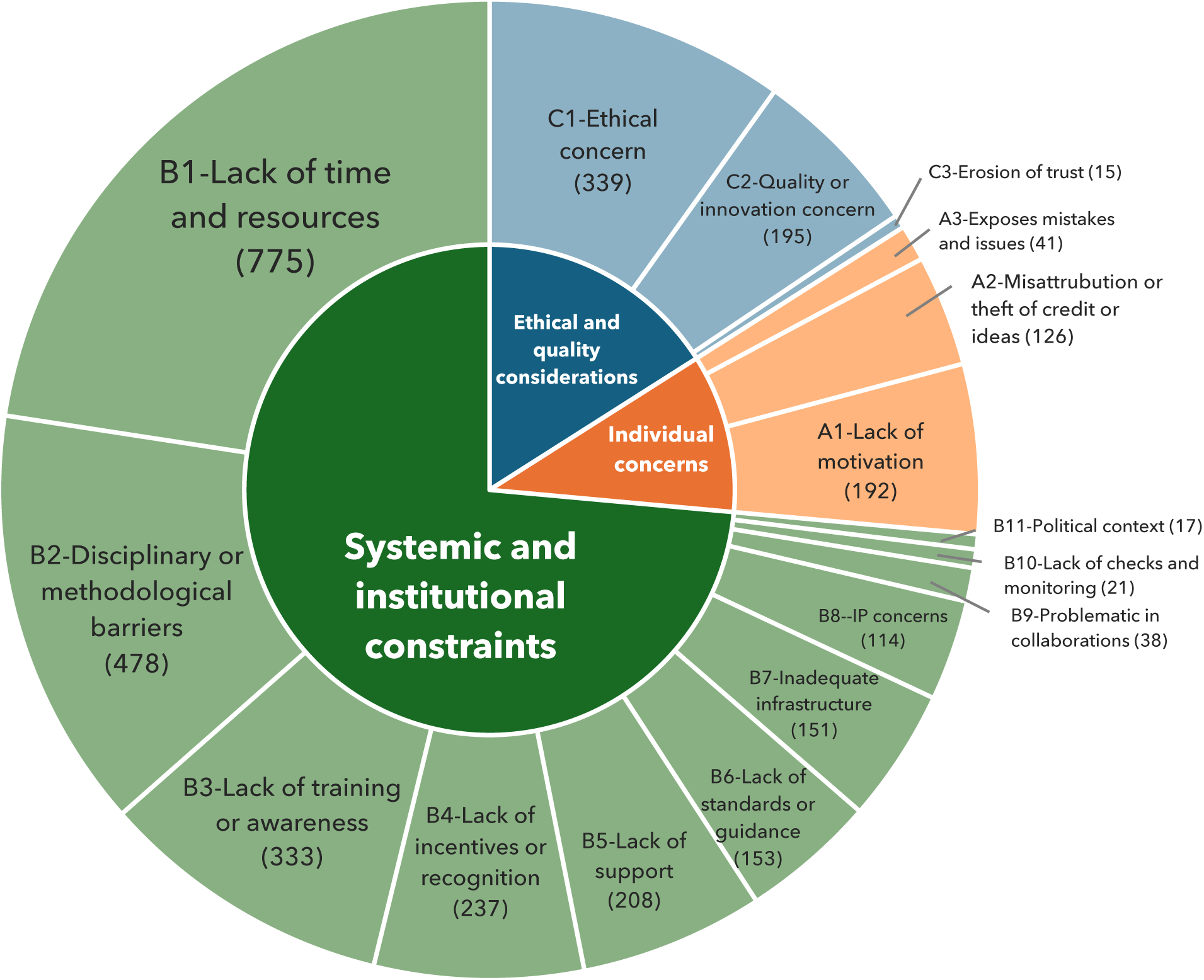
Frequency of themes and clusters of risks and barriers to open and responsible research practices.

#### 3.2.1 Theme A1 - Lack of motivation

An important theme mentioned in 192 comments throughout the survey was the lack of motivation to undertake open research practices. This theme was mostly mentioned as an issue respondents noticed from colleagues and from the community, rather than from themselves. Comments such as *“People in the field don’t care and don’t have time to care”* or *“Even though I think it is good science my team have no interest in pre-registration”*. Several references to open research not being integrated in the culture of research were also mentioned. In addition to these remarks, some respondents indicated that they themselves lacked motivation to undertake such practices, for example by saying *“not worth effort”*, *“seems unnecessary work”*, *“waste of time and leads to duplication”*, or *“If it was enforced it would waste a lot of my time to no good purpose”*. Respondents described part of this lack of motivation as resulting from the perception that open research practices were merely *“a tick box exercise”* or that they have no real impact and lead to additional work for sharing elements that are unlikely to be reused.

For some more specific open research practices such as creation of Open Source Software, version control, and computational reproducibility, respondents also expressed that such practices should not rely on researchers, and instead require support from institutions and qualified staff. Illustrative quotes include *“This is important, but impractical if the burden falls entirely on the researcher”*, *“It is not what I am employed to do”*, or *“I’m not a computer scientist, I moved away from that field on purpose because I found it wasn’t for me. It would be nice to have a group of comp sci people to work with to make our own programs, I think it’s silly to expect biological scientists to not only be expert scientists, but also writers, figure illustrators, coders, mentors, public speakers, and teachers.”*

Opposing this lack of motivation however, at least 287 counter-nodes ascertained the importance of open research practices, for example by qualifying the practices as *“very important”*, *“essential”*, or *“crucial”* for good research, with one participant even mentioning that a specific open research practice was “hugely important to all humanity” (i.e., in this case computational reproducibility). Given that the question only asked about risks and barriers and not about the importance of open research practices, the high density of answers in this line is notable.

#### 3.2.2 Theme A2 - Fear of theft of credit

Another recurrent theme which was mentioned in 126 references throughout the survey relates to a fear of *“being scooped”* and losing one’s *“competitive advantage”*. This was mentioned mostly in relation to preprints or pre-registration, but was also visible in other open research practices such as open codes and open methods.

> *“There are concerns about disclosing techniques, code etc. that took enormous personal investment to develop-these can then be easily used by others, reducing your professional advantage.”*

In this theme, respondents expressed a worry that bigger groups with more resources may be able to conduct experiments or analyses from the shared material faster than those sharing the material and may take the credit and originality away. For example, one respondent mentioned that they *“would worry that groups with larger resources could take these ideas and improve upon them/finish work quicker.”*, with others mentioning that this is an even greater risk in *“’hot’ topics.”*. One respondent summarised this idea by stressing that *“[c]ollecting data involves a vast amount of effort. Giving it away to people who haven’t put that effort in doesn’t always feel like a smart career move.”*

Related to the above, several respondents also mentioned that the practice of using guidelines for recognising the specific substantive contribution of everyone involved in research projects (topic ix in the survey), could cause or exacerbate conflicts within teams and even lead to delays in the publication process.

> *“It sometimes leads to misunderstandings and minor issues as some people do not feel recognised enough.”*
>
> *“There are always worries that if you don’t acknowledge/include someone (despite no substantive contribution from them), they will feel insulted or spiteful.”*
>
> *“A recent example where this has provided a barrier is on a conflict for correct authorship credit delaying publication significantly. The delay was because a postdoc was being bullied by their PI and the department did not have an answer for how to enforce the fairest outcome.”*

#### 3.2.3 Theme A3 - Fear of exposing mistakes

Finally, some respondents were also concerned that open research practices could expose errors or issues with their work and make them vulnerable to criticism (41 references). This was reflected in respondents expressing worries in sharing their data, codes, or other research material because they feared that these open outputs could expose errors, untidy practices, or other issues such as animal rights concerns, even *“in the most well-intentioned research plans and methodologies”*. Respondents mentioned, for example, that *“the stress of including a silly mistake or typo in one’s open analysis code is the biggest barrier (for me)”*, or that *“There is always a concern someone may attack you over your data or methods”*.

### 3.3 Cluster B - Systemic and institutional constraints

Numerous systemic and institutional constraints that posed a barrier on the adoption of open research were also mentioned. This cluster was the biggest overall, including 2,525 references that ranged from lack of time and resources to commit to open research, methodological or disciplinary barriers, or infrastructural barriers. Respondents also mentioned that institutions did not support them adequately, did not provide appropriate training, and did not recognise, incentivise, or monitor open research practices, making it difficult for them to engage in open research.

#### 3.3.1 Theme B1 - Lack of time and resources

The most mentioned barrier highlighted that a general lack of time and resources was a tangible barrier to open research practices. This theme was captured in in 775 references, making it the most mentioned barrier overall. Most participants mentioned that open research takes time and that they found difficult to engage in open research practices, with several suggesting that they were *“[d]rowning in busy work already”*.

In addition to general comments on the time needed to undertake open research practices, almost 60 responses insisted on the additional administrative burden that open research practices – or the institutional requirements necessary to enable them – would generate. Comments such as *“Yes, waste of time with bureaucracy!”* or *“the paperwork involved is prohibitory”* illustrate these concerns. In addition, over a hundred comments discussed administrative burden or challenges from ethical approval or data protection processes which prevented open research practices. For example, some mentioned that *“ethical clearance needed for fieldwork usually takes too long”* or that *“Data management plans can be too restrictive.”* Many of the comments in this area discussed the delays caused by the needed approvals, and one respondent even described that for them, the *“enormous”* delays *“have put the viability of the relevant projects in doubt”*. In other instances, it was the amount of training and learning needed to overcome the complexity of some open research practices that prevented adoption, still referring to research staff being stretched for time.

> *“I have done GDPR training about four times in the last 24 months due to mandatory requirements in different third level educational institutions. The level of time spent doing mandatory training on these things leads to a measurable reduction in research time available. This is a barrier.”*
>
> *“The ability to share code is extremely important, but it can be an inefficient use of time to learn new coding just so it is for open source software.”*

Finally 154 references mentioned that the cost of open research practices was a barrier in itself. This largely referred to Article Processing Charges in the question about Open Access (142 of the 154 references), yet other types of costs were also mentioned, such as costs needed for research co-production. Related to this last point, several responses also mentioned that limits in funding structures prevented open research practices. For example, several comments suggested that *“you are just being paid to publish the work, not spend time afterwards making it accessible”*. Other comments, mostly related to co-production, mentioned the opposite effect, missing time prior to grant funding to establish efficient partnerships.

> *“Designing a new study in a co-production way requires funding to engage with the community before applying for larger research grants. These funds are not explicitly advertised or not widely available.”*
>
> *“Ethical partnerships are difficult to accomplish with charitable and community stakeholders because our projects are fixed-term and grant/output-driven, so building up long-term relationships can be very difficult”*

#### 3.3.2 Theme B2 - Disciplinary and methodological barriers

In many instances, participants mentioned that the open research practices mentioned were not applicable to their research or discipline. This was the case in 478 comments, with responses such as *“not relevant to my field”* or *“This is not applicable to my research”*. In many of such cases, the lack of relevance was pointed as relating to open research practices being perceived as inadequate for qualitative or non-empirical research. For example, one respondent highlighted the following: *“In the humanities, it is important that findings are different and shed a different light on the ‘data’. Demanding reproducibility is missing the point entirely. If this is enforced, it will damage the discipline.”* Another, while discussing transparent qualitative data, mentioned: *“Again, not directly a concern for non-empirical researchers. Although it is often assumed to be - usually by those with little understanding of what philosophers actually do!”* These responses highlight a disciplinary gap in how open research practices are understood and in their perceived relevance for different types of research and methodologies. Several respondents also worried about the fixed nature of pre-registrations, supporting that their research requires inductive and evolving approaches. This was often raised by qualitative researchers, but also raised as an issue for discovery science, as well as longitudinal, observational and exploratory research. A few comments indicating this are shown below.

> *“Using a pre-determined way of analysing qualitative data is problematic, e.g., the collected data may not fit the assumptions that a certain method of analysis will be optimal (e.g., it may not be detailed enough for IPA, but reflexive thematic analysis would offer new insights). Hypotheses are not used in qualitative research. Taking this approach would also undermine inductive data analysis.”*
>
> *“I do not agree with fixing all parameters before conducting the research - some of my most impactful publications have been from finding phenomena in the data which I was not looking for”*.
>
> *“Prescriptive processes introduce risk of becoming ideas development rather than research.”*
>
> *“I do not feel this is appropriate for my (non-experimental) field of research. It risks closing off too many fruitful avenues and serendipitous research findings.”*
>
> *“Yes, people need to be trained in knowing what data looks like, but the thing is anything is potentially data. And you have to be able to bluff, in a project proposal and a grant application, and say you know what kind of data you are going to look for and etc. But the best research projects find so much more than they think they will, and often of an entirely different project/question. So the whole performance of proposals and hypothesis are a dance. What we need is a system that trains scholars to be ethical subjects in the midst of research, and then sets them free to study whatever happens to be in front of them.”*

In line with these disciplinary and methodological barriers, a few respondents worried that the increasing expectations of open research practices *“diminishes and devalues observational and non hypothesis driven studies”*, limiting the *“range of possible studies that are considered ‘good’”* and building *“a very narrow view of what good research is.”*

#### 3.3.3 Theme B3 - Lack of training and awareness

Lack of training and awareness was mentioned as a barrier to open and responsible practices, captured in 333 references. Most comments mentioned that more training would be needed or that training should be more visible and accessible to all. While most comments pushed for *“more training”*, several comments also stressed that the training offered should be improved. In fact, several respondents mentioned that current training was *“too basic”*, *“generic”*, or even *“low grade”* and *“unappetising”*, and that efforts should be invested to make the training more adapted to researchers’ needs. We also coded lack of awareness with the open and responsible research practice in this theme, with almost half (143) of these 333 references pointing to a lack of awareness or understanding about the open research practice mentioned. Comments such as *“Barriers are mainly lack of knowledge around new ways of working.”* or *“people do not know what ‘data’ is and assume this does not apply to them”* expressed a perception that a lack of awareness was a barrier to the practices, but several respondents also expressed that they themselves did not understand what the practice implied, for example saying *“I have no idea what this is”* or *“not heard of this before.”*

That being said, at least 43 counter-nodes suggested that training was already sufficiently provided or that it would not be needed. For example, someone responded that *“[t]raining is clear and research governance staff very supporting”*, while another responded that *“I don’t feel you need training on this. You just do it.”*. Some of these comments mentioned that the real challenge was to move the practices to become the norm.

Comments such as “Does not need training - just making the norm” and “Again, training not the issue so much as enforcement and compliance” express this point. Other comments suggested that respondents felt able to teach themselves: “I train myself. I don’t need courses. I have a phd which is a license to learn”.

#### 3.3.4 Theme B4 - Lack of incentives or recognition

Another important theme, mentioned 237 times, was the lack of incentives and recognition for open and responsible practices, a theme which unavoidably leaked into a lack of motivation for undertaking the practices mentioned (theme A1). In this theme, comments recognised that open and responsible practices take time and efforts but are rarely recognised in career advancement. This theme was especially present when discussing open access – where respondents feared that focusing on open access publications would conflict with the incentive to publish in high impact journals since in certain disciplines *“top journals are seldom open access”*. Similar concerns were phrased with regards to the publication of preprints, where several respondents explained that preprinting their work may *“[p]reclude publication in certain journals”* which were relevant for advancing their career. Other practices were also mentioned as conflicting with current recognition and reward models, potentially going against career progression and be ‘*‘perceived to impact competitiveness”*. Additional comments point to a genuine worry that open research practices may become a risk to researchers’ careers if formal recognition for such practices is not targeted.

> *“[T]he institution uses messaging to encourage such work but there is no reward or recognition so it is seen as a risky activity in terms of desire for progress and career success.”*
>
> *“[T]his [preregistration] would be murder for any early career scholar. Shocking that some disciplines do this.”*

Interestingly, 10 counter-nodes hinted at the opposite, stating that open research practices were important for advancing one’s career. For example, one response mentioned that it is *“important for early career researchers to learn how to do this competently in order to progress their career.”*. A few of these comments mentioned that Open Access in particular was important for the Research Excellence Framework (REF), others mentioned that knowing how to manage data or enable computationally reproducible analysis was important in preventing issues during inspections and audits, and others mentioned that practices such as Open Access and preprints could help generate higher visibility and chances of citation.

#### 3.3.5 Theme B5 - Lack of support from institutions

Related to the previous themes, 208 responses indicated that a major barrier to embracing open and responsible practices was the lack of support from their research institution. Some descriptions of the university as being *“risk adverse”*, providing *“little encouragement”* or adopting a *“passive approach”* showcase this perceived lack of support. In many cases, participants mentioned that the practices are *“recommended but not required”* and that the uptake of the practice ultimately *“relies on individual researcher initiative”*, without formal support.

In the same order of ideas, a few respondents also pointed to a lack of capacity or understanding from the institution, sometimes mentioning that the practice would require the university to provide competent staff with expertise in these practices to support adoption, but that these were either absent, *“unhelpful”*, or *“good but overstretched”*.

Finally, some respondents suggested that beyond the organisation as a whole, senior staff were not engaging in the practice and therefore unable to mentor and support the practice within their research groups. Some quotes below illustrate this issue.

> *“[O]lder staff members often aren’t on board with reproducibility practices.”*
>
> *“My field is collecting larger and larger quantities of data. Most lab heads do not have experience with data management plans.”*
>
> *“Help would have been appreciated but just not on the horizon of my supervisors.”*

It is worth mentioning however that 29 counter-nodes also mentioned that open research was sufficiently supported by research institutions, with several comments pointing specifically to *“[e]xcellent support”* provided by their libraries.

#### 3.3.6 Theme B6 - Lack of standards or guidance

Related to the above, we identified 153 comments which mentioned that the lack of guidance or the lack of standards make it difficult to adhere to open research practices. These included comments on the lack of guidance to understand how a practice should be undertaken, as well as a lack of guidance and standards within universities on their preferred approaches. Comments such as:

> *“It is very frequently mentioned without really clear steps for how to achieve it.”*
>
> *“[T]here are different standards of ‘reproducible’ though, so easy for someone to get lost down a rabbit-hole”*.
>
> *“The University fails to make these standards part of its own research guidelines”*.

The guidance for some of the open research practices was described as *“diverse and scattered”*, *“not always clear”*, and *“inconsistent”*. In some cases, respondents also mentioned that *“there are different conventions”* between disciplines, projects, or even geopolitical contexts (e.g., regarding GDPR regulations) which complicate adoption of open research practices. Finally, several respondents mentioned that open research standards are rapidly evolving.

> *“Keeping up to date with the latest requirements and mandates can be a challenge.”*

As a result, barriers expressed in this theme suggest that for some, the lack of clear standards on what is expected with regards to open research practices is a barrier to adoption, making these practices *“hard to follow”* and *“confusing”*.

Interestingly, 76 counter-nodes mentioned that open and responsible research practices are either required from universities, journals or funders, or that *“additional centralised rules or training”* is not needed or could even have a negative impact. Another 114 counter-nodes mentioned that open research practices were already done as a standard practice, also suggesting that further standards may not be universally desired.

#### 3.3.7 Theme B7 - Inadequate infrastructures

Another systemic barrier mentioned as blocking adoption of open and responsible research practice was the inadequacy or incompatibility of infrastructures currently used in research (151 references). This theme was particularly prominent in discussions of open source software, defining data and code, version control, and computational reproducibility.

In this theme, respondents often mentioned that the software or infrastructure generally used in their fields were not compatible with open research standards, but that moving away from these usual infrastructures was difficult. For instance, someone mentioned that *“The standard software used for qualitative research tends to be proprietary (and better supported by universities)”* and that taking the time and effort to learn open source options was deterring, leaving them no viable option other than to use the usual software. In some cases, proprietary software was seen as essential for successful publication or large-scale influence:

> *“It can be harder to get published if you chose not to use particular bits of proprietary software.”*
>
> *“Software currently used for my field of study are proprietary and all our existing IP, analysis, models, research … relies on these software. Changing to open source options would be preferable but risks destroying the networks of influence that we have built within the UK over time.”*

Some responses also worried about the long-term interoperability of open source alternatives, mentioning a *“[r]isk of future researchers not being able to access [their] work”*. One respondent explained that *“Open source software is rarely supported over a long enough timescale, and is often poorly documented and sometimes badly written”* while another mentioned that *“open-source software usually is less mature, more volatile with a higher initial learning curve.”* In some cases, respondents also pointed to universities limiting access to open source software for security reasons.

Finally, several comments mentioned that current infrastructures were inadequate for the type of research they were working on. This was particularly mentioned for large data sets and the lack of large storage capacities from universities or open sharing platforms.

#### 3.3.8 Theme B8 - Intellectual property concerns

Concerns around conflicts between intellectual property rights and open research was visible in 114 comments throughout the survey. Comments coded in this theme referred to the difficulty of sharing data, codes and methods when collaborating with industry, or when hoping to build commercial interest from the research findings.

> *“Major barriers! I can’t patent a drug (and so it becomes impossible to develop it) if it is in the public domain.”*
>
> *“It is difficult to release open source when working with stakeholders who have existing non-open-source license requirements.”*
>
> *“Any code that I produce would be proprietary under the terms of my funding.”*

Comments in this theme also mentioned issues with the original accessibility and openness of the data used. In fact, several respondents mentioned that the data they used was only accessible upon request or offered only limited access, making it difficult for them to share research based on such data and making open research practices which required providing access to others (e.g., co-production) challenging. In addition, reusing data whose ownership was external was also mentioned as an issue, for example in data sharing.

> *“Private ownership of data places a barrier to sharing data and code.”*
>
> *“I primarily work with sensitive and national health/education/crime datasets which cannot be shared (i.e. cannot make open data) as I am not the data owner.”*
>
> Issues with copyrights and licensing were also noted, with several respondents mentioning that the rules around these were *“hard to navigate”*.
>
> *“I don’t know what’s okay to share sometimes and what copyright to use.” “Understanding which Open Source licence to use when where has taxed us a little.”*

#### 3.3.9 Theme B9 - Problematic in collaborations

A slightly smaller theme raised issues with collaborations when conducting open and responsible research (38 references). Respondents here mentioned that different adherence to, and views on open research practices between collaborators could create friction and challenge adoption.

> *“Although I think open research is important for its own sake, it throws up certain challenges/frictions when working with others who do not share this view. The lack of particularly strong recognition/reward or compliance requirements from my institution means they do not feel they need to bother with taking on my perspective.”*
>
> *“[W]ith international collaborations it can be hard to agree to make code open if not all collaboration members agree.”*

While the comments above mostly highlight different levels of interest and desire to uptake open research practices, other comments raised concrete barriers which could impact collaborations or be impacted from large collaborations. This was the case for version control, which was mentioned by a few respondents as difficult to uptake in large teams despite being even more important within those. Unequal access to software was also mentioned as a potential issue, with at least one comment mentioning that *“collaborators abroad may not have access to the same software as we do in the UK and so sharing data / analysis may be challenging”*.

#### 3.3.10 Theme B10 - Lack of checks and monitoring

Several comments also mentioned the lack of checks and monitoring impacting how open and responsible research practices are taken up. Although this theme was only mentioned in 21 comments, it raised an important challenge in which the lack of enforcement of open and responsible research practices meant compliance was lacking. Comments mentioned that these practices were *“not well policed”* or that *“[t]here is no stick”* for inadequate practices, meaning that bad practices continue and open practices keep a low uptake. Beyond the practices themselves, respondents also mentioned that compliance with the planning of the research was not monitored: *“In order to get ethical approval I need a DMP for my work; no-one checks I comply with it though.”*. Finally, some respondents even worried that their institution may turn a blind eye to lack of compliance for reputational benefits:

> *“COI [Conflict of Interest] again involves politics – supervisors have vested interests in companies or their own financial interests which are not dealt with well – the university turns a blind eye to this and if you raise this, they don’t want to know because often those supervisors are people with a lot of power.”*

#### 3.3.11 Theme B10 - Political context

The last theme captured in the cluster of systemic and institutional constraints was the political context acting as a barrier to open research. This was a minor theme that included only 17 comments. Of these, the majority were repeated comments from the same respondents which included *“Brexit is destroying the UK”*, *“Brexit is destroying UK higher education”*, and *“Brexit impedes everything we do”*. While these comments were not entirely relevant nor explicitly related to open and responsible research practices, a minority of comments were more directed at the issue. Some examples included mentions about halted publications since the war in Ukraine – *“The war in Ukraine has meant that ATLAS has stopped publishing papers while the issue of what to do with Russian institutions is resolved.”* – changes to Medical Device Regulations since Brexit, or issues participating in Erasmus programmes which was seen as a barrier to open research.

### 3.4 Cluster C - Ethical and quality

On an ethical and quality level, respondents worried that open research practices introduced new ethical concerns, or even that they detracted from high-quality research and innovation. Ultimately, some comments worried that open and responsible research practices may dampen the trust from the public in science. This cluster was found in 549 comments.

#### 3.4.1 Theme C1 - Ethical concerns

The most mentioned theme in this cluster was the worry that open research practices may conflict with ethical and confidentiality standards, especially when working with sensitive populations and restricted topics (339 comments). Respondents particularly mentioned a risk to participant confidentiality when sharing certain types of data such as qualitative data, longitudinal data, local or small-scale research data, data from research with minority groups, or video data, as well as research in certain disciplines (e.g., anthropology where participants may *“reveal extensive biographical information”*).

Respondents also mentioned that informing participants that they would make their data open may deter participation, making participants *“reluctant to take part if anonymised transcripts were made available in a repository”*, or *“reluctant to be recorded”*, for example. Some respondents even mentioned that, when participating in research, they themselves would be reluctant to share their data if they were given the choice: *“As a participant I never consent to my qualitative data being shared beyond the research team.”*

Restrictions needed to enable anonymisation were also thought to potentially impact the research, for example by requiring to exclude vulnerable people and minorities, or by shaping how participants react and respond. Respondents worried that even where participants are willing to share their data, the mere awareness that their data will be shared may have *“epistemology and positioning”* impacts, with participants risking to be *“less comfortable sharing on sensitive topics if they knew full accounts would be made available.”* Similarly, some respondents worried that curating the data to enable sharing may require it to be *”‘cleaned’ to the point uselessness”*. One respondent explained that *“Interviews are very context-rich and to redact all identifying information would make them quite thin as well as possibly change the meanings of the responses.”*

Beyond the risks to research participants and research, respondents also mentioned that in some cases, risks of dual use or security concerns prevented them from embracing open research practices. Some respondents, for example, explained that their work in the nuclear sector or in defence was subject to strict export controls and accessibility requirements, sometimes limiting the collaborations they are allowed to build.

Other respondents mentioned that opening their data to non-expert could be dangerous and potentially *“pose a risk of misinterpretation of data by non-professional”*. Concerns that data may be used *“for purposes outside of it’s ethical approval”* were mentioned by a few respondents. This barrier was mostly mentioned with regards to data sharing, but other practices were also targeted; for example, one respondent expressed concern that *“using open source software”* may open their work to a larger pool of analysts which *“may allow people less qualified in aspects unrelated to the software to get in the mix”*.

Risks relating to research integrity and questionable research practices were also mentioned in several instances. While most of these comments mentioned a *“risk of non-compliance”* with a relevant open research practice (thus not raising a risk from the open practice per se), a few comments also explained that open research practice did not prevent failures of research integrity. For instance, a few respondents mentioned that *“pre-registering a protocol does not prevent an alternative protocol being used in practice”*, that *“irresponsible authorship practices are common”* despite contributions being acknowledged openly, or that *“[c]onflicts of interest are often overtly apparent in publications where none are declared”*.

Another type of ethical issue that was seen as a barrier to open research was linked to the abuse of for-profit players benefiting from open research expectations, especially open access requirements. Participants commented that *“publishers are gaming the system to their own advantage”*, and that publication costs are *“a terrible investment of funder money to be tipping it into the pockets of for-profit publishers.”*.

In rare cases, responses worried that open and transparent research practices risked being altogether detrimental to research and the academic system. Some examples include:

> *“Transparency is impossible. In fact it is detrimental. It puts people at risk. There have to be safeguards against transparency. It is good practice to show how un-transparent one is, methodologically, by indicating how you anonymise, etc. The academic industry is being run into the ground by people flouting nonsense terms without a grasp of the danger this presents to people who participate in our research.”*
>
> *“HUGE problems. FAIR is unethical, when you come down to it, as our data is either a legal/ethical land field or ‘cleaned’ to the point uselessness. funding bodies require the use of repositories, but the repository archivists (thankfully) have a better head than the funders.”*
>
> *“OA [Open Access] is yet another example of the colonial attitude of the sciences. The consequences of OA have not been thought through by UKRI etc in regards, for example, to the ecology of small learned societies.”*
>
> *“We always collaborate in research, and we know how to collaborate and share, but a bunch of parasitic science bureaucrats try to formalise the ‘open science’ agenda as a money/career-making vehicle for their bureaucratic machinery. ‘First invent the obstacle then offer your help”’*

These more extreme views often connected to the next theme of open research potentially reducing the quality of the research being performed.

Finally, a few responses, particularly in response to the question on open access, hinted at a risk of inequalities being accentuated between individuals, research groups, countries, and disciplines from open research.

> *“Open Access is a trap. It serves to separate the rich from the poor while claiming to do the opposite.”*
>
> *“It [Open Access] affects REF returnability of publications and not all publications are fiscally supported by the institution to be open access. It therefore creates academic inequity within the institution.”*
>
> *“People with generous funding face almost no barriers, people without access to funding face massive barriers.”*

#### 3.4.2 Theme C2 - Quality concerns

The second theme we identified in some of the respondents’ comments related to a concern that open research practices may reduce the quality or value of research (195 references). Some comments from previous themes already hinted at this element, with respondents worrying about the value of open research practices or the impact that open research practices may have on the integrity of the data. In this theme, some practices were more targeted than others. In particular, co-production (41 comments), open software (34 comments for use and 10 for creation), and preprints (27 comments) were often targeted.

Regarding co-production, comments expressed a number of concerns, including that co-production *“undermines researcher objectivity”*, that it may skew results, or that it *“reduced the expert nature of research in some areas”*. Some mentioned that, when done wrong, co-production can become tokenistic and as such, it *“can have damaging effects to individual community members as well as the field itself (causing lack of community/public trust in science).”* Other comments were more directly opposing co-production to quality, stating that the focus on co-production may act as a distraction from excellent, innovative science, with some even believing it may reduce the competitiveness of the country’s research and innovation as a whole.

> *“Funding applications that require evidence of co-production is a barrier to focusing on good science.”*
>
> *“Research that practitioners and stakeholders etc. are interested in co-producing is not always at the cutting-edge of academic debates.”*
>
> *“The emphasis is and should be on outstanding science not on politically correct mediocre science that involves stakeholders who are not scientists and who slow down our national efforts to be world competitive. We are trying to compete with China, with the US, with Europe.”*
>
> *“Why (and how!?) would patients or other non-research trained stakeholders be involved in biomedical research? We are trained (to PhD) level to understand and carry out research. How do you propose we involve the public? This sounds like another nonsensical initiative that will waste our time and resources.”*

Issues relating to quality were also often mentioned in the topic of open source software. Most of these comments simply referred to respondents who mentioned that open source software could be *“poorly documented and sometimes badly written”* and that it was sometimes *“less mature, more volatile”*. This was linked to a lack of standards (e.g., ISO standard) in this area.

Many comments, including very strong opinions, were also raised about preprints. The comments mentioned that since preprints are not peer-reviewed, they potentially allow immature and *“untested data”* into the public domain. For example, one respondent mentioned that preprints introduced *“a danger of false messages getting out”* referring to issues that arose during Covid-19. Another respondent went a bit further, stating that *“Preprints allow unreviewed garbage to be published”*, while another linked preprints to a risk that unethical research may become public given the lack of peer-review.

> *“I’m concerned about how preprints allow for unethical research to be disseminated with the veneer of science. I’ve seen racist scientific work particularly picked up at preprints and shared widely before peer review can address it (and peer review doesn’t do this well). Also, they really are not a thing in many fields, including mine.”*

Along the same lines, respondents mentioned that preprints should only be used for *“a manuscript that is in a finished state, ready to be submitted to a journal.”*, but many comments suggested this was not always the case.

Other responses also opposed open and responsible research to innovation, stating that it could stifle originality, *“hinder blue-sky thinking”*, and lead to a *“[r]educed ability to innovate”*. Possible delays in publications caused by open research practices were also mentioned, alongside a risk that the quality of journals selected may be impacted by open access requirements – as mentioned also in the theme on lack of incentives and recognition.

#### 3.4.3 Theme C3 - Erosion of trust

Interwoven with the above two themes, 15 comments raised a concern about the overall trust in science and the impact that open research practices may have. While this theme was much smaller than other themes and raised very different perspectives, we found the theme important to mention on its own. Trust went both ways. For example, a lack of trust from partners was mentioned as a barrier to efficient co-production, but as seen above, poorly executed co-production was also mentioned as a risk to public trust in science.

**Fig 2.**
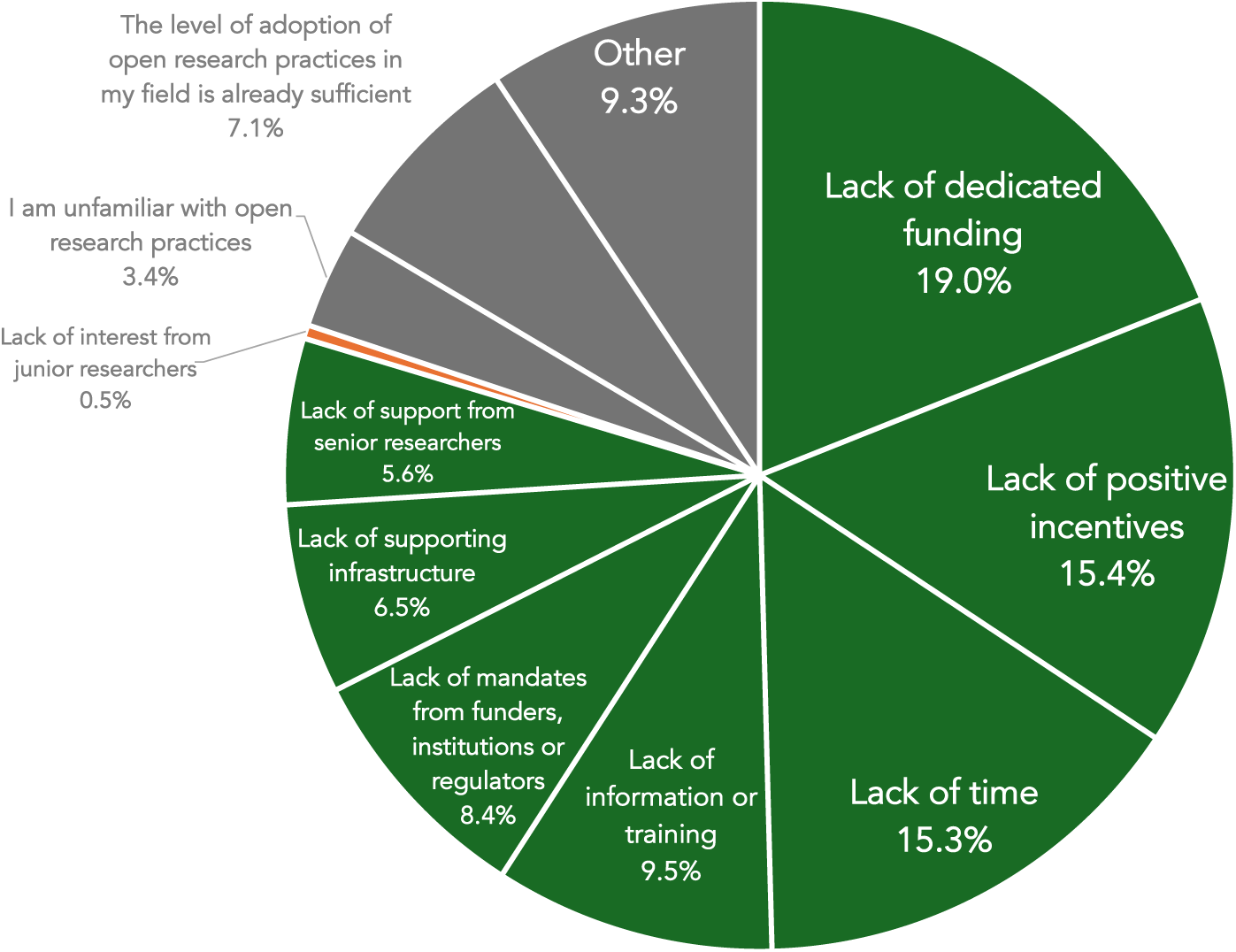
Distribution of responses on the final multiple-choice question from the survey which asked respondents to select *“the greatest barrier to the uptake of open research practices”* in their field.

The importance of trusting researchers and not basing trust only on open practices was also mentioned, pointing to a risk that *“research can be stymied by over strict, one size fits all regulations.”* A couple of responses pointed to a lack of trust in universities and academic systems as a whole, raising concerns that go beyond open research practices.

> *“I don’t trust the system and so I try to be cautious and think with my students as much as possible outside the formal systems since the barriers can be poorly calibrated so they prevent ethical work and enable unethical work”*.

However, from the comments mentioned in the counter-nodes, at least 3 comments mentioned open and responsible research practices as a condition for trust in science, and 3 comments mentioned it as necessary for the credibility of research.

#### 3.4.4 Question 16 - Greatest barrier

The last question of the survey, although separate from the above analysis, is also relevant to look into since it asked respondents *‘Having considered barriers above, which do you perceive to be as the greatest barrier to the uptake of open research practices in your field?’* Respondents were offered a choice between 11 possible responses, as we will see below. A total of 1,081 respondents provided answers to this question. See Figure 2 for a distribution of findings.

While these responses were not seen by the qualitative analysis coder (NAB) prior to the analysis to preserve the inductive approach to the coding, themes identified in the qualitative analysis resulted in a close enough match from the answer choices provided in this last question to enable a vague comparison. Interestingly, responses from participants to this last question somewhat mirror the density of responses captured in the inductively identified themes through the qualitative analysis.

While the qualitative analysis of open-text questions found ‘Lack of time and resources’ to be the most mentioned theme (775 answers, or 22.6% of answers), the multiple-choice question identified ‘Lack of dedicated funding’ as its top barrier (19% of responses), and ‘Lack of time’ as its third biggest barrier (15.3% of responses). The second biggest barrier identified by the final question was ‘Lack of positive incentives’ (15.4% of responses). This barrier was less prominent in the qualitative analysis, but was still in the top 5 concerns with 237 responses (6.9% of mentioned barriers and risks). ‘Lack of information or training’ came fourth in the multiple-choice final question with 9.5% of responses, which is very similar to what was found in the qualitative responses where 333 answers (9.7%) pointed to a ‘Lack of training or awareness’. The multiple-choice questions’ next biggest barriers were ‘Lack of mandates from funders, institutions or regulators’ selected by 8.4% of respondents, ‘Lack of supporting infrastructure (e.g., sufficient storage for open data / publishing platform for open monographs)’ selected by 6.5% of respondents, and ‘Lack of support from senior researchers (e.g., supervisors and principal investigators)’ selected by 5.6% of respondents. These could correspond to qualitative analysis themes of ‘Lack of standards and guidance’, mentioned in 153 comments or 4.5% of mentioned barriers and risks, ‘Inadequate infrastructure’ mentioned in 151 comments or 4.4% of barriers and risks, and ‘Lack of support from institution’ mentioned by 208 comments or 6.1% of mentioned barriers. Finally, the survey final question obtained 0.5% (N=5) of responses selecting a ‘Lack of interest from junior researchers’ as the biggest barrier. This barrier has no equivalent in the qualitative analysis. Similarly, several barriers found in the qualitative analysis have no equivalents in the multiple-choice provided. The pre-defined choices almost exclusively addressed systemic and institutional barriers, with little to no consideration to individual or ethical and quality concerns. This gap reinforces the added value of the inductive process in identifying additional barriers that may not have been considered otherwise.

Beyond these choices, the final question also obtained 7.1% of respondents stating that *‘The level of adoption of open research practices in my field is already sufficient’* (similar to counter-nodes in the qualitative analysis), 3.4% of respondents declaring *‘I am unfamiliar with open research practices’*, and 9.3% (101 respondents) selecting *‘Other. Please explain’*.

A quick qualitative overview of the 101 responses mentioned by those selecting ‘Other’ shows 74 responses relating to the cluster of ‘Systemic and Institutional Barriers’, 13 responses relating to ‘Individual barriers’, and 8 responses relating to ‘Ethical and quality concerns’ (note, responses were sometimes placed in multiple themes and clusters, explaining the higher count from adding all cluster counts).

What was especially interesting in these other answers however, was responses which suggested that the risks and barriers of open and responsible research practices are profoundly interconnected, and that trying to solve only ‘the biggest’ barrier would probably not make a viable change. We found 16 answers that suggested this interconnection, some of those examples are detailed below.

> *“There is not one single ‘most important’ barrier there are multiple mutually-reinforcing barriers.” “Many of the above. Selecting just one option is not possible as they are all interlinked.”*
>
> *“Genuinely all of the above - big multivariate problems don’t have a single “biggest” contributor. As in - if you only fix one of the above, nothing will change.”*.
>
> *“I think this oversimplifies the issue. There is not one single ‘most important’ barrier there are multiple mutually-reinforcing barriers. Most are mentioned above, however based on open access experience mandation [mandated] by funders without provision of resources, time and support could be positively harmful to research culture and practice.”*

## 4 Discussion

This study examined perceived risks and barriers associated with 14 open research practices across 15 UK HEIs. Our findings show that difficulties adopting these practices cannot be reduced to a lack of awareness or willingness across the research community. Instead, barriers operated across three interrelated clusters: individual concerns; systemic and institutional constraints; and ethical and quality considerations. Lack of time and resources was the most recurrent theme, but it was closely connected to insufficient training, inadequate infrastructure, unclear guidance, limited institutional support, and a lack of recognition within existing career and reward structures. Respondents also identified substantive risks, including loss of intellectual credit, exposure to criticism, conflicts with intellectual property, threats to confidentiality, inappropriate application across disciplines, and possible adverse effects on research quality. Taken together, the findings suggest that uptake depends not simply on persuading individual researchers of the value of openness, but on whether research environments make responsible openness feasible, appropriate and professionally sustainable.

### 4.1 Interconnected Barriers to Adoption

Across respondents, open research was frequently described as something that must be undertaken in addition to already demanding research, teaching, and administrative workloads. This points to a broader structural issue: institutions and funders increasingly expect open research practices to be implemented across the research lifecycle, yet the ones conducting and enabling the research do not always perceive that they have been provided with the necessary time, resources, infrastructure, or support required to embed open research effectively. This notion is reinforced by the pattern of barriers identified throughout the survey. Time constraints, funding limitations, inadequate infrastructure, lack of training, insufficient guidance, and weak institutional support emerge not as independent obstacles, but as indicators of a common challenge: open and responsible research is not yet deeply embedded into the wider research landscape. Adoption therefore remains dependent on individual effort and personal commitment. This aligns with a growing body of literature suggesting that cultural change in research cannot be achieved solely through awareness raising and policy mandates [17, 18], but requires institutional investment and systemic alignment of incentives, resources, and expectations.

Our findings also highlight tensions between open and responsible research practices and the lived realities of members of the research community working in existing systems. Respondents frequently questioned whether the considerable effort required to produce openly accessible outputs, share date and code, or engage in other related practices is recognised for career advancement and research funding. This suggests barriers to adoption may arise when researchers are asked to work according to one set of values while continuing to be assessed according to another set of rules. Efforts to increase adoption are therefore likely to be most effective when open research practices are recognised and rewarded as legitimate scholarly contributions rather than additional activities undertaken alongside *“core”* academic work or as a nice to have. Similarly, funding systems that not only require open research practices, but also embed the time and resources for these practices within the funding structure are needed to ensure that all involved are able to adopt open research practices without disadvantage.

Another important consideration is the diversity of research traditions represented in the survey data. Respondents often questioned whether Open Research practices were relevant to research in the humanities and qualitative, observational, or exploratory research. This epistemic discussion reflects assumptions originating in the Open Science movement and experimental fields. These concerns demonstrate that debates surrounding open research are not simply about adoption versus resistance. Rather, they reflect ongoing discussions about what transparency, reproducibility, and rigour should mean across different epistemological traditions. The challenge for institutions and funding bodies may therefore be less about promoting universal adoption of specific practices and more about broader goals of openness, FAIRness, and responsibility while respecting methodological diversity.

### 4.2 Perceived Risks of Open Research

An important finding of the survey study is that many concerns raised by respondents reflected tensions between open research practices and other responsibilities that researchers are expected to uphold, including protection of participants, ensuring research quality, maintaining professional credibility, securing Intellectual Property.

This distinction is important because discussions of open research often frame adoption as a choice between openness and reluctance to change. The present findings however suggest a more nuanced reality.

Respondents frequently appeared supportive of the principles underpinning open research, yet questioned how these principles should be balanced against competing ethical, professional, and methodological obligations. In this sense, the risks identified in the survey may be understood less as barriers to cultural change and more as expressions of the trade-offs researchers perceive when attempting to implement openness in practice.

Concerns relating to participant confidentiality, intellectual ownership, professional recognition, and research quality suggest that respondents do not view openness as an absolute good, but as one jigsaw piece that must be balanced against competing ethical, methodological, and career considerations. This finding supports a shift from open research as unrestricted openness and transparency towards a model of responsible research that recognises legitimate constraints on sharing and is sensitive to epistemological considerations that open research may have. Importantly, concerns about being scooped, exposing mistakes, or reducing research quality are unlikely to be resolved through advocacy alone, as they are rooted in the incentive structures, competitive pressures, and accountability mechanisms that continue to shape academic research. Whether these perceived risks are justified is less important than the fact that they influence researchers’ willingness to engage with open research. The challenge for institutions and funders is therefore not simply to promote openness, but to demonstrate how openness can be implemented in ways that safeguard academic rigour, recognise contributions, maintain confidence, and mitigate against the risks identified in the survey.

### 4.3 Towards Responsible Research Implementation

Taken together, these findings suggest that the debate surrounding open research may need to move beyond the question of whether the research community supports openness or not. Most concerns identified in this study did not challenge the value of transparency, accessibility, or reproducibility. Instead, respondents mostly questioned how these principles can be implemented responsibly within practical realities of contemporary research.

Therefore, we might want to consider a shift from promoting open research practices *“in principle”* towards supporting its implementation in practice. This will likely require investment in infrastructure, specialist support and training, alongside reforms to incentive and career progression structures. However, it will also require recognition that openness is not a uniform objective. Successful implementation will depend upon maintaining sufficient flexibility to accommodate diverse disciplinary requirements and methodological approaches.

The challenge therefore is not primarily a negative attitude towards open research, but whether the institutions, the wider system, and research environments enable it.

### 4.4 Limitations

The survey captured an important snapshot of open and responsible research practices in UK HEIs. However, a few issues are important to note. First, the rate of completion of the survey impacts how the results can be interpreted. As mentioned above, 2,567 participants took part in the survey. Of these, 1,221 (48%) answered at least half of the survey questions, including 1,081 (42%) who completed the survey. Consequently, given the structure of the survey which addressed 13 questions for each of the 14 open and reproducible research practice mentioned in the order described in the methods, practices mentioned near the end of the survey will have reached fewer respondents than practices mentioned in the beginning of the survey. Quantitative comparisons between the different open research practices is therefore not advised.

The survey also attracted several (148) comments raising frustrations with the survey. Most of these comments related to the length of the survey which took substantially longer than expected for respondents. A few comments also suggested that some respondents did not understanding what the question asked (responses generally classified as ‘unclear or irrelevant’), that some respondents were frustrated about the comment box requiring a response despite the question framing it as optional, or that they did not understand what the open research practice definition meant (i.e., survey text related to open research practices was hyperlinked to the FORRT glossary, see the data descriptor for further details [12]). While these responses highlighted important issues, we consider that genuine answers still contained highly valuable information and deserved to be analysed given the high commitment of those who took the time to provide detailed and thoughtful answers. The critical comments were used carefully to inform the design of the next UKRN survey which was circulated in 2025-2026.

Finally, given the qualitative nature of this review, certain interpretations and thematic classification of comments could have been interpreted differently by a different coder. NAB did the coding, but revisions of uncertainties were discussed with LH-N and the overall themes emerging from the comments were discussed with LH-N and AJS. Revision of codes, clusters, and themes, and revisions in the coding of specific comment therefore took place throughout the analysis process. To promote transparency on the coding and to enable reanalyses, a new dataset has been created which includes all comments extracted as containing relevant risks or barriers, and the themes, clusters, and open research practice they belong to. This dataset is available on Figshare ^1^ [16].

## 5 Conclusion

Respondents broadly recognised the value of open and responsible research but identified both substantial barriers to adoption and important risks associated with implementation. While barriers largely reflected structural challenges related to resources, incentives, and institutional support, perceived risks centred on professional vulnerability, ethical responsibility, and research quality. Together, these findings suggest that increasing uptake requires more than encouraging researchers to be open. It requires creating research environments in which openness is properly supported, appropriately recognised, and implemented in ways that remain sensitive to diverse research contexts and responsibilities.

## Acknowledgments

We would like to thank everyone at the UK Reproducibility Network (UKRN) who supported the OTRP Survey and all participants for their valuable time and input.

## Footnotes

1 https://doi.org/10.48420/33287682

## Notes

https://doi.org/10.48420/33287682

### Competing Interest Statement

The authors have no conflict of interest to declare. The authors are however members of the UK Reproducibility Network. AJS is Institutional Lead for The University of Manchester, LH-N is Local Network Lead for The University of Manchester, and NAB is Open Research Coordinator and Administrator for The University of Manchester.

